# Keystone competitor creates spatial patterns of biodiversity

**DOI:** 10.1101/2023.04.21.536496

**Authors:** Katherine K. Ennis, Ivette Perfecto, John Vandermeer

**Affiliations:** Integrative Biology Department, University of California, Berkeley, Berkeley, CA 94720; School of Environment and Sustainability, University of Michigan, Ann Arbor, MI 48109; Department of Ecology and Evolutionary Biology, University of Michigan, Ann Arbor, MI 48109

**Author notes:** Corresponding author: Katherine Ennis, Department of Integrative Biology, University of California, Berkeley, Berkeley, CA 94720. Katherine K. Ennis –. Ivette Perfecto –. John Vandermeer –.

**Keywords:** Competition, coexistence, keystone species, spatial patterns, ant, richness

## Abstract

Conventional ecological theory on competition and competitive exclusion states that competition should limit diversity. However, diversity of all species is more common than competitive exclusion would suggest, especially in the tropics. Here, we use theoretical and field approaches to examine the relationship between a keystone ant competitor (*Azteca sericeasur*) and the richness of a ground-foraging ant community in a spatially explicit context. Theoretically, we demonstrate that a fixed keystone competitor can increase species diversity. In addition, we sampled the ground-foraging ant community in three differentially managed coffee habitats and found – with the exception of plots in the most intensified coffee habitat – a consistently higher species richness near to the keystone competitor and lower richness with increasing distance. These patterns show that keystone competitors may contribute to the maintenance of local species richness.

## 1. Introduction

Species co-existence and the maintenance of species diversity has been a focal topic in ecology for decades. Many studies, both theoretical and empirical have demonstrated the importance of factors like habitat complexity, disturbance, predation and competition. Among the many more specific ideas explaining species co-existence is the notion of a keystone species. First demonstrated with the starfish *Pisaster ochraceus* in the rocky intertidal zone of the Pacific coast of North America [1], keystone predation has since been demonstrated widely across many terrestrial and aquatic systems [2-4]. Application of the keystone concept has expanded beyond to non-predatory species describing interactions other than predation. For example, several studies identify keystone mutualistic interactions[5-7] while others have demonstrated that a resource can act as the keystone species in a community [8, 9]. Yet another study [10] identifies a keystone intimidator species; a species whose effect on the community stems from modifying the behaviour of another species, or a non-consumptive effect.

There are two related components to the keystone mechanism. First, competition may vary from weak to strong [11]. If competition is weak, a predictable equilibrium of species densities is expected. If competition is strong, however, competitive exclusion permits only one species can survive in a particular niche [12]. Several recent studies have highlighted the inherent paradox of communities that are thought to exhibit both strong competition and simultaneously high levels of species diversity [13-15]. In theory, if competition structures communities then competitive exclusion should limit shared use of ecological niches and community richness should be equal to the number of niches. However, many communities have such high species richness that habitats with an equivalent number of unique niches seem unlikely. Alternative explanations posed to explain this paradox suggest that founding effects and variable outcomes of competition due to environmental variability may be more important than strong competition resulting in competitive exclusion [13]. The latter explanation reveals how competition may still be important to communities but not in the traditional competitive exclusion and niche partitioning sense [15].

In light of these ideas the competitive process may explicitly determine the surviving species, or the outcome may be indeterminate, and founding densities of each species generally determine which species will survive. Although these scenarios represent ends on a continuum, some communities near the strong competition end of the spectrum do exist. Communities experiencing strong competition are particularly common when space is the limiting resource. Such strong competition is commonly believed to structure ant communities [16].

Second, the “interruption” of competition in a strong competition context can dramatically alter the outcome of the competitive process. Notable examples of this process include a keystone predator that consumes competitively dominant prey [1, 17], the intermediate disturbance hypothesis that predicts that the process of completive exclusion by one species is in effect “reset” by a disturbance [18, 19] and the disruption of dominance in competition hierarchies due to behavioural responses to parasitism [20-25]. In these examples, what would normally be a reduction to one (or a few) species through strong competition is interrupted such that the expected competitive reduction to one species is repeatedly delayed and the observed number of species remains, on average, high.

### (a) A theoretical framework

These ideas come together in a particularly interesting way when cast in a spatial context. Suppose the competition intensity for a suite of *S* species is uniformly large (in the Lotka-Volterra context suppose *α* > 1.0), but there is another species, call it the keystone species, which has an even greater competitive effect on all the others, but is fixed at some point in space. As a result this keystone species imposes a gradient on the community such that it moves from a state of strong “interruption” to a state of no “interruption” with increasing distance from the keystone species. We propose that under these conditions, which are likely common in nature, there will be a gradient in the competitive assembly of species ranging from classical indeterminate competitive exclusion to control by a specific dominant competitor (where the keystone is dominant), and that between those two competitive structures, the indeterminate versus dominant control may effectively cancel one another, producing a local maximum in species diversity.

Formally, we present a model that corresponds to this basic framework, assuming the effect of the keystone species is given as *β*, is,

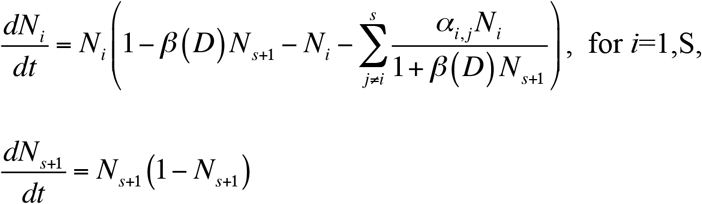

where *N*_*i*_ is the biomass or density of the *i*th species, *α*_*i,j*_ is the competitive effect of species *j* on species *i*, and *β* is the indirect competitive effect of the keystone species on all the others, and is a function of *D*, the distance from the fixed position of the keystone species. Clearly, if species *s*+1 (the keystone) has no access to the competitive arena (i.e., where *β*L*D*L-> 0), the species that happens to have the advantage at the beginning of a competitive bout will completely take over the community, whereas if species *s*+1 has access (i.e., *β*L*D*L >>0), there will be a point at which the keystone reduces the competitive abilities of the other species sufficiently such that only the keystone will survive. Between those two extremes we might expect the permanent or semi-permanent survival of all the species in the community. The value of *β*, thus represents a position on the postulated interruption gradient. Analytically equation set 1 produces intuitively reasonable results (if *β* -> 0, one species dominates through competitive exclusion, as *β* becomes large only species (s+1) survives, and with intermediate values of *β*, the effective competition coefficients of the system (*α*_*i,j*_/(1+ *β*(*D*)*N*_*s+1*_) become small (but *β* is not sufficiently large to allow *s*+1 to dominate) thus leading to coexistence of the other species, and finally as *β* becomes very large, it completely dominates the community and the keystone species is the only survivor.

Using this basic framework, we constructed a spatial model that initiated a random group of species on a spatial lattice and fixed a purported keystone species in the centre of the lattice. Using the above equations, we then calculated the resulting number of species at cells located at various distances from the position of the keystone over time (see Appendix S1). An exemplary result is displayed in figure 1, reflecting the qualitative generalization that there should be a zone of higher species diversity between two zones of lower diversity. Thus, equation system 1 suggests a generalization that when a keystone competitor is restricted, as it might be if located permanently at particular points in space, its effects on an indeterminate competition community may produce a pattern in which a zone of high species diversity is sandwiched between two low species diversity areas (figure 1).

**Figure 1.**
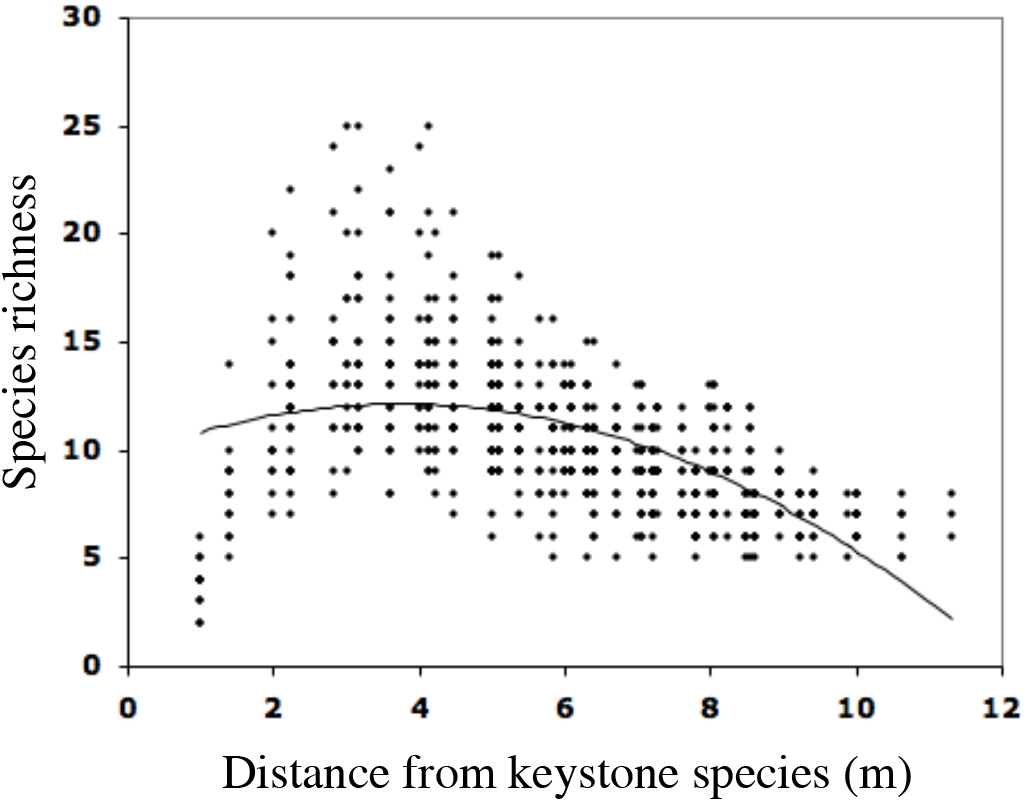
Simulations of species diversity as a function of distance to keystone species, based on Equations 1 (see data supplement, Appendix S1, for full explanation).

Here we report on a field study investigating the role of a particularly aggressive ant species, *Azteca sericeasur*, in affecting the spatial pattern of species diversity of a ground foraging ant community in general, across three different coffee agroecosystems in southern Mexico. The system is particularly apt for testing the above ideas. The ground-foraging ant community in this region is diverse and relatively well known, and exists in an evident non-random spatial pattern, sometimes qualitatively resembling the mosaic pattern so frequently reported for ants in agricultural systems [26, 27] where multiple dominant species may form distinct, mutually exclusive territories.

We suspect that this pattern is largely determined by competitive interactions among the species, a common assumption for ground-foraging ants [16, 28, 29]. Embedded in this system is the purported keystone species, *A. sericeasur*, a canopy dominant ant that nests exclusively in arboreal sites, but is found foraging on the ground in the immediate vicinity of its nest. Thus, the ground is covered with mosaics of generalist foraging ants and in a fixed position is the keystone. We then test the prediction that the pattern of this community should broadly resemble the computer simulations presented in figure 1.

## 2. Material & Methods

### (a) Study Site

The study was conducted in two adjacent coffee farms, Finca Irlanda and Finca Hamburgo, in the Soconusco region of Chiapas, Mexico during the rainy season (May – August). The sites are approximately 40 km NE of Tapachula (15° 11’ N, 92° 20’ W). The habitat of the coffee agroecosystem in general is relatively homogenous compared to neighbouring natural habitats due to the prevalence of the planted coffee and associated shade trees. Finca Irlanda is a 280-ha. shaded organic coffee farm with a uniform distribution of shade trees that represent over 100 shade tree species [30] and encompasses two management styles referred to in this study as high and moderate shade coffee production. Within the active production area, shade cover is moderate with between 50-70% shade (moderate shade). In this production area we have established a 45-ha. plot with approximately 11,000 shade trees mapped and surveyed for presence of *A. sericeasur* annually for the past 6 years [31]. The high shade area of Finca Irlanda produces coffee, but is virtually unmanaged and has approximately 70% canopy cover (high shade). In this area we have established another 6-ha. plot using the same methods for tree marking, location and hosting *A. sericeasur* data. Finca Hamburgo is a large (1000 ha.) coffee farm that has much lower shade cover (15-30%), much lower shade tree diversity and employs the use of more conventional management techniques including the application of fungicides, herbicides and occasionally, insecticides. While there is no established plot in Finca Hamburgo we have located all nests of *A. sericeasur* in a 10-ha region of the farm. Following a standard coffee production classification system [32], the high shade area is consistent with the description of a traditional polyculture, the moderate shade area with a commercial polyculture, and the low shade area with a shade monoculture where total canopy cover is lower and most shade trees are conspecifics or congenerics.

*Azteca sericeasur* is a dominant canopy nesting ant found throughout the Neotropics in coffee agroecosystems [30]. It forms large carton nests in trees, but can be found foraging on the ground, in the leaf litter and in coffee. It is a well-documented mutualist of the hemipteran coffee pest, the green scale insect, *Coccus viridus* [33]. *Azteca sericeasur* is further associated with many organisms including parasitoids, other hemipteran species, fungi, beetles, spiders, birds and ants [34]. Many of these interactions with the broader invertebrate community are a result of both direct interactions and indirect interactions driven by chemical pheromone cues from *A. sericeasur* [35, 36]. Annual censuses of *A. sericeasur* throughout the 45-ha plot reveal that at the landscape level *A. sericeasur* nests form a clustered distribution and occupy only 3% of all shade trees [37]. This suggests that *A. sericeasur* is not a numerically abundant ant species relative to other ant species found within the plot. The accumulated study of the interactions between *A. sericeasur* combined with a general understanding of it’s relative distribution and abundance across the landscape makes *A. sericeasur* a likely keystone species of this habitat [34]. The associated ground-foraging ant community is made up of many species from the *Solenopsis* and *Pheidole* genera but species of *Gnamptogenys, Pachycondyla, Acromyrmex, Paratrechina, Brachymyrmex* and *Odontomachus* are also common.

### (b) Ant sampling

Ant surveys took place in June and July in the morning hours between 7:00-11:00 am at each of the three coffee production areas (low, moderate, and high shade) within eight plots. In the high shade we established two plots (plot H1: 24m × 20m; plot H2: 20m × 20m), in the moderate shade we established four plots (plot M1: 48m × 48m; plot M2: 40m × 40m; plot M3: 32m × 24m; plot M4: 24m × 24m) and in the low shade we established two plots (plot L1: 24m × 24m; plot L2: 24m × 24m). Each plot was established around a cluster of *A. sericeasur* nests residing in shade trees above the coffee bushes. The variability in plot size results from various idiosyncrasies in the local topography and vegetation. The number of *A. sericeasur* nests in each plot varied from 2-8. For all plots we established a grid with a sampling point either at every 4m or at every 2m (depending on the size of the plot) across the entire area of the plot. We used gridded plots rather than transects because the clustered distribution of *A. sericeasur* nests made it difficult to identify areas that lay outside the influence of *A. sericeasur*. Furthermore, a grid design also provided more robust sampling to the frequently variable ant richness found at the bait scale.

We baited for ground ants at all 8 sites using tuna in oil, thus defining our community as ground-foraging ants that are attracted to baits. At the two largest sites (M1 and M2) we placed one teaspoon (5g) of tuna on the ground every 4 m forming a grid pattern, and at each of the remaining smaller sites we placed one teaspoon (5g) of tuna on the ground every 2m. We waited no more than 20 minutes and recorded each species found within 10 cm of the bait. Species occurrences at baits can be dominated by strong recruiting ant species and limit the observed occurrences of weaker species. However, small-scale time series of bait occupancy reveal that strong recruiter species require approximately 20 minutes to completely exclude and deter other species (e.g. [38]). Based on this understanding, we proceeded to check baits within 20 minutes of placement so that, on average, we assessed the baits prior to exclusion of weaker species by strong recruiter species. For individuals we could not identify consistently to species or morphospecies in the field we collected a sample for identification in the lab using the guide developed by Jack Longino on the *Ants of Costa Rica*. For species level identification of the most commonly occurring species we confirmed the identification with Dr. Jack Longino.

### (c) Statistical analyses

Richness of species was calculated for each lattice point within each plot. We summed the observed number of different species at the point in question plus the 8 nearest neighbour points surrounding the bait (the Moore neighbourhood). This was done to quantify those species that were in the area but were not observed near the baits. In the analysis, we eliminated those baits on the edges because the richness of those points could only be informed by 5 neighbour points for the edges and 3 neighbour points for the corners. We then regressed this neighbourhood richness (where each point/bait is equal to the number of unique species in the focal bait and the 8 neighbour baits) of ground-foraging ant species against the distance to the nearest nest of *A. sericeasur* and chose the best regression model with both linear and quadratic models, comparing the sum-of-squares of the two models with an “extra sum-of-squares” F test [39]. This test is similar to the Akaike Information Criterion (AIC) in that it incorporates the goodness-of-fit (as measured by sum-of-squares) with the number of parameters used in the model. For each site, we compared the linear and quadratic models according to this equation:

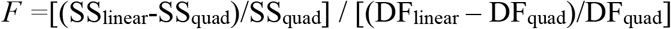

Where SS_linear_ is the sum-of-squares of the residuals in the linear model, SS_quad_ is the sum-of-squares of the residuals in the quadratic model, and DF is the degrees of freedom of the linear and quadratic models where indicated. The test is based on the idea that if the null hypothesis were true then there would be relatively little difference between the difference in the sum-of-squares and difference in the degrees of freedom [39].

## 3. Results

Across all habitats ants were found at 94% (1203/1282) of baited points. Occupancy was similar across all three habitats –89% occupancy in high shade, 96% occupancy in moderate shade and 96% occupied baits in low shade. *Pheidole protensa, Pheidole synanthropica, Solenopsis geminata* and a *Pheidole* sp.1 were the most common species found on the baits across all farms. While the overall species composition varied considerably between plots, the most common species were similar across all coffee habitats. Plot H2, however, had the most distinctive composition as only one of the most commonly encountered species listed above (*Pheidole* sp. 1) was encountered in H2.

In both high shade plots (H1 and H2) we found significant negative relationships between species richness of ground-foraging ants and the distance to the nearest nest of *A. sericeasur* (figure 3a, b; Plot H1: *r*^*2*^=0.196, *F*_2,96_*=*11.8, *P* < 0.0001; H2: *r*^*2*^=0.288, *F*_2,78_ *=*15.74, *P* < 0.0001). This relationship was significantly stronger using a quadratic model rather than a linear model (Plot H1: *F*_97,96_ *=* 23.33, *P* <0.0001; H2: *F*_79,78_=13.41, *P* <0.0001).

In the moderate shade plots M1, M3 and M4 we found significant negative relationships between species richness and distance to the nearest *A. sericeasur* nest (figure 3c,e,f; Plot M1: *r*^*2*^=0.290, *F*_2,118_ *=*24.14, *P* < 0.0001; M3: *r*^*2*^= 0.049, *F*_1,119_*=*6.12, *P* = 0.015; M4: *r*^*2*^=0.133, *F*_2,162_*=*12.49, *P* < 0.0001). The relationship was not significant in Plot M2 (figure 3d; *r*^*2*^ *=* 0.039, *F*_2,78_*=*1.6, *P* = 0.208), but still showed significant improvement in the relationship with a negative quadratic equation, as did Plots M1 and M4 (M1: *F*_119,118_ *=*38.91, *P* < 0.0001; M2: *F*_79,78_=2.52, *P* <0.0001; M4: *F*_163,162_=22.58, *P* <0.0001). Plot M3 did follow the same negative trend as the other plots with moderate shade but the added explanatory power from the quadratic regression was not significant (*F*_79,78_ = 1.08, *P* = 0.33).

In the low shade Plot L2, there was a positive relationship between species richness and distance to the nearest *A. sericeasur* nest, however that relationship was not statistically significant (figure 3h; L2: *r*^*2*^ = 0.032, *F*_1,119_*=*1.19, *P* = 0.091). However, in Plot L1 there was a significant positive, non-linear relationship found in species richness and increasing distance from *A. sericeasur* (figure 3g; L1: *r*^*2*^=0.283, *F*_2,118_*=*10.02, *P* < 0.0001). Only Plot L1 in the low shade coffee habitat benefitted significantly from using a quadratic model (L1: *F*_119,118_ = 10.02, *P* < 0.0001; L2: *F*_119,118_ = 1.18, *P* = 0.117).

## 4. Discussion

The keystone species concept has been widely applied to numerous systems [1-10]. Our study advances the keystone concept by applying it to a behaviourally dominant competitor using first a theoretical model and then an empirical study in a spatially explicit context. The findings from both perspectives demonstrate that this keystone competitor may play a significant role in maintaining species diversity in communities. Prior studies in coffee agroecosystems illustrate the extent of behavioural and non-consumptive effects that *A. sericeasur* has on various components of the insect community [24, 40-42]. However, the effect of *A. sericeasur* on the surrounding ground-foraging ant community is not limited to alterations in ant behaviour. Communities of sessile organisms or similarly, central place foragers (e.g. ants) have long thought to be structured by strong competitive interactions, especially those communities in agroecosystems [43]. Here, the keystone interaction between *A. sericeasur* and the species that comprise the ground-foraging ant community is a function of both aggressive behaviour and resource competition with species of the same taxonomic assemblage.

We demonstrate that an aggressive, behaviourally dominant keystone species may act as a deterrent to the inevitable competitive exclusion resulting from strong competition between members of the ground-foraging ant community. The resultant spatial pattern reflects this mechanism both theoretically (figure 1) and empirically (figure 3a-c,d,f). This idea is similar to the ideas expressed in tri-tiered competitive hierarchies [14], but the details of hierarchical theory have not previously applied to elucidate spatial pattern, nor has it been thought to apply to a single species that both constitutes a small relative abundance and is behaviourally dominant. In this case the aggressive arboreal nesting ant, *A. sericeasur*, is associated with changes in the spatial pattern of species richness of the ground-foraging ant community in coffee agroecosystems. This relationship is complex and varies with shade cover, but the general trend forms a “halo” pattern (e.g. figure 2). That is, *A. sericeasur* appears to have a negative effect on species richness very close to its nest, a positive effect at a distance between 5-10m, but with increasing distance, species richness declines again (figure 3a-e), as would be expected in areas of strong competition from competitive exclusion.

**Figure 2.**
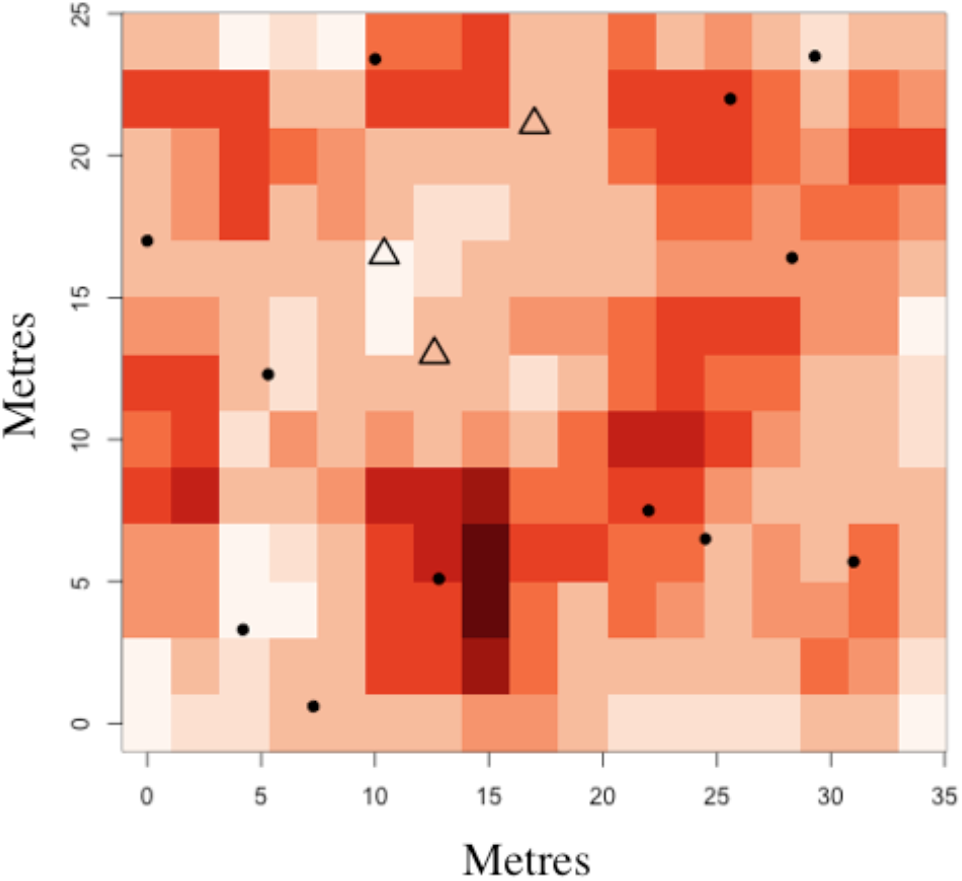
An example of the graphical representation of ant richness results found at each bait in for one moderate shade plot, M3 (32 × 24 m). Points represent shade trees, triangles represent shade trees with a nest of *A. instabilis* and darker colour squares indicate more species. (See data supplement, Appendix S2, for a graphical depiction of the results from each plot).

The obvious exception to this pattern occurs in the two plots in the low shade coffee agroecosystems. In these plots, the relationship between species richness and distance from *A. sericeasur* nests is inconclusive: it may be an example of a breakdown of the general negative trend, or it may be that the relationship is actually reversed (figure 3f,g). These results suggest that these interactions within a habitat are altered by habitat disturbance. As *A. sericeasur* is primarily an arboreal-foraging ant, changes in management intensity that lead to lower canopy resources may direct the foraging behaviour of *A. sericeasur* towards seeking resources on the ground. This would increase the contact that ground-foraging ant species have with *A. sericeasur* and could have an impact on the competitive relationships themselves. It is also possible that low shade coffee has altered the scale at which a negative and non-linear relationship is visible. For example, it may be that all ant species in the low shade habitats have greater foraging distances because there are fewer available resources and thus the scale of the plots were simply too small to capture the full pattern.

**Figure 3.**
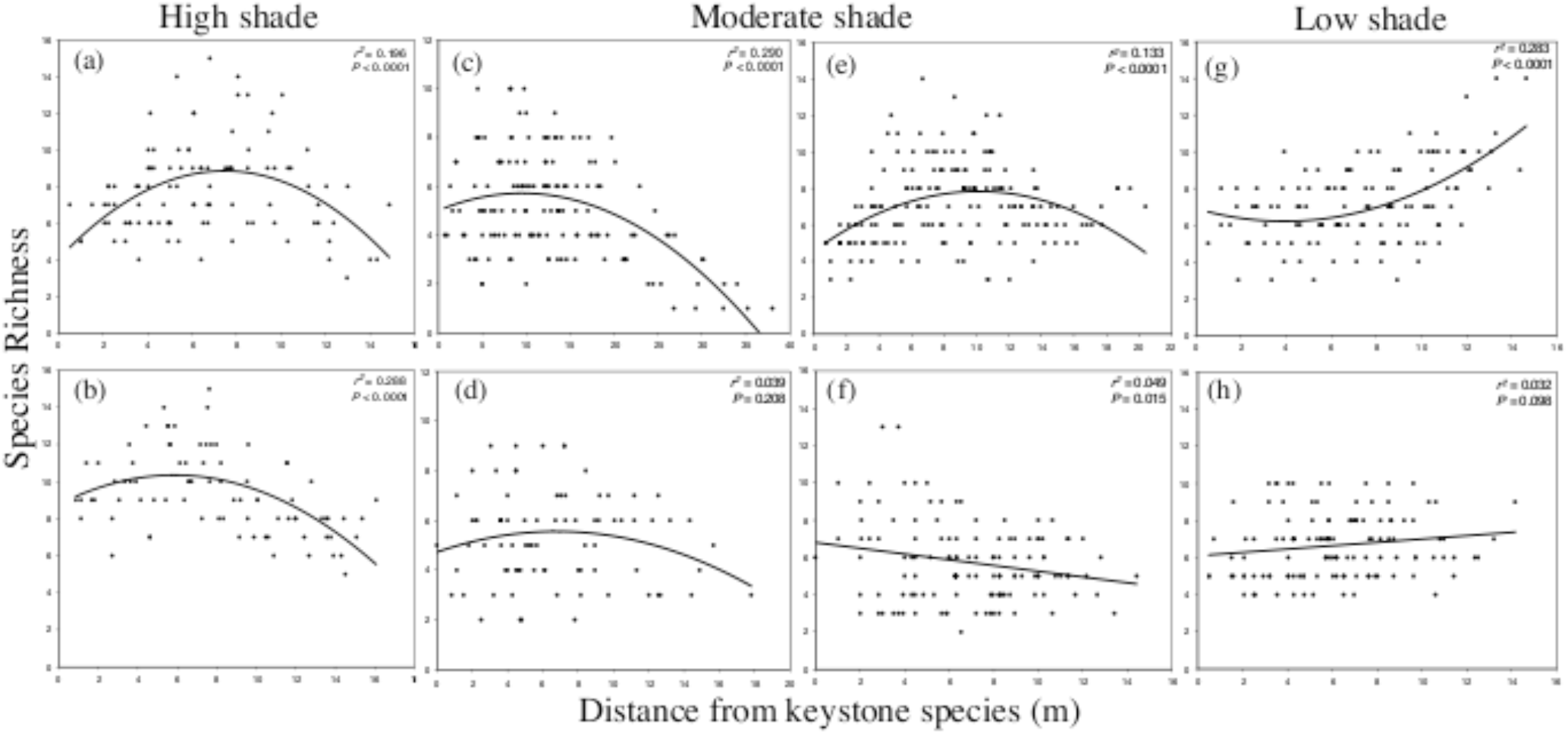
Regressions of species richness against the distance to the keystone species (nearest *A. instabilis* nest) for each site, organized by shade quantity. Graphs (a, b) represents high shade Plots H1 and H2; (c-f) represents moderate shade Plots M1-4; and (g, h) represent low shade Plots L1 and L2, respectively.

Our results are broadly consistent with previous studies showing that dominant species can affect the structure of local communities, although our empirical observations of the details of spatial pattern have not been emphasized before. Spatially explicit theoretical models demonstrate that the positive effect of keystone predation on species diversity requires consistently intermediate resource productivity [44-46]. Other spatially explicit models for ant communities suggest that the spatial pattern we find both theoretically and empirically is dependent on a competitive balance between the interacting species [47].

Empirical studies of dominant ant species have previously demonstrated their effect on ant community structure. In twig-nesting ant assemblages, *A. sericeasur* affects nest colonization processes of the most common arboreal twig-nesting species, but did not affect more rare species [48]. Co-occurrence patterns from ground-foraging ant assemblages in Australia have been previously associated dominant ground-foraging ant species are associated with higher species richness [14]. Likewise, experimental removal of a single dominant species in a ground-foraging ant assemblage found that the exclusion of a single dominant species increases the dominance of other ecologically similar species, but did not result in increases in species richness overall [49]. In addition, a moderate abundance of multiple dominant ant species has been shown to promote higher species richness [50]. However, in all studies, the dominant ant(s) were numerically dominant, rather than exclusively behaviourally dominant as with *A. sericeasur*. In the first example, the three patchily distributed dominant species (*Iridomyrmex* sp., *Papyrius* sp. and *Oecophylla smaragdina*) are associated with overall greater richness. In this case, the various dominant species span both ground (i.e. *Iridomyrmex* sp. and *Papyrius* sp.) and arboreal ant (i.e. *Oecophylla smaragdina*) assemblages and make up a large portion (30-77%) of ant abundance per site. In the latter two cases, however, the studies examine dominant ant species that are found within a single ant assemblage and are numerically dominant species. In the second example [49], the dominant ant, *Iridomyrmex purpureus*, makes up between 68-84% of all ants present, while in the third example [50], the five dominant ant species make up a combined 54.5-72.2% of all ants present. In contrast, *A. sericeasur* is found on fewer than 4% of all baits and often makes up < 2% of all ant occurrences. Thus, our study demonstrates that species that are behaviourally and competitively dominant, yet in effect numerically rare, may still play a major role in determining the spatial structure of ant communities.

We initially identified the theoretical basis for the effect of a keystone competitor in the context of strong competition through space and then found empirical support in the field for the findings from the model. Within the lesser-disturbed habitats (those that displayed negative trends between richness and distance from the keystone species), we suggest that the increase in species richness near the *A. sericeasur* nests could be because the keystone competitor disrupts the strong competition between ground-foraging ants. A similar explanation has been proposed as part of the revised ‘interstitial hypothesis’ where the increase in richness associated with dominant ant species is a result of an indirect positive effect between dominant and subordinate species [14]. The positive effect conferred on subordinate species occurs by way of neutralizing the negative effects of the subdominant species on subordinate species. Clearly the trends found are not entirely in keeping with the proposed theory, suggesting that the interactions present in the coffee habitats are influenced by disturbance and in particular by management intensity. Our study builds on the specific understanding of the behaviour of *A. sericeasur* and more generally on the competition that exists in ant communities to demonstrate in a spatially explicit context how through both competition and intimidation (behavioural, non-consumptive effects) a keystone species contributes to the number of species found in a local habitat.

## Acknowledgements

We thank A. Andersen and an anonymous reviewer for helpful comments that significantly improved earlier versions of this manuscript. We also thank D. Allen for his important statistical and graphics help, D. Gonthier and S. Philpott for insightful comments and support, B. Chilel and O.G. Lopez-Bautista for field assistance, and J. Longino for his help in identification of specimens.

## Data accessibility

All datasets may be located upon publication in the Dryad repository.

## Funding statement

Funding provided by NSF DEB 0349388 to I. Perfecto and J. Vandermeer, and NSF DEB 1020096 to

S. Philpott.

## Appendices

### Appendix S1

#### Model specifications and derivations

Representing, theoretically, the idea of a keystone competitor could take a variety of forms. If the intended keystone simply entered the equations as just another competitor in the LV sense, it could have no effect that would be any different than any others, other than perhaps be a super competitor, as,

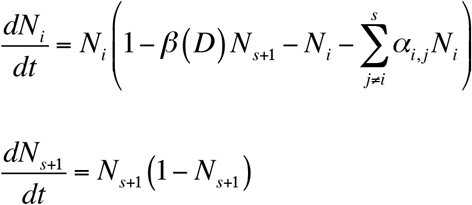

in which case it would simply outcompete everyone else, when *D* is low.

Equations 1 were formulated in a spatial context on a 15 × 15 lattice, more-or-less corresponding to the size of our plots in the field. At each lattice point we presumed that the density of each species was the sum of its densities at that point plus the eight surrounding points (the Moore neighbourhood). Thus, equations 1 were solved with Euler’s simple method, employing the logarithmic form with Δ*t* = 1, namely,

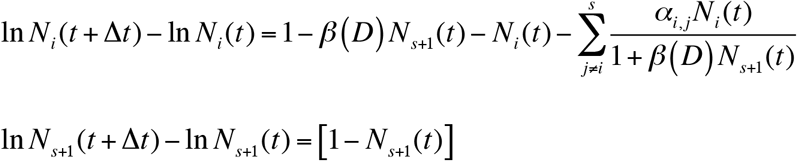

We allow β to be related to D as;

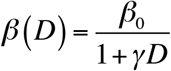

where *β*_0_ is the base line competitive effect of the keystone (when *D* =0), and *γ* is the rate at which that effect declines with distance from the fixed position of the keystone. The model was run four times with parameters set as *β*_0_ = 10^3^, *γ* = 40, and competition coefficients were chosen from a random uniform distribution ranging from 0 – 3 (i.e. the mean competition coefficient was 1.5). The initial number of species was 50 and the model ran for 20 iterations each run.

### Appendix S2

**Appendix S2.**
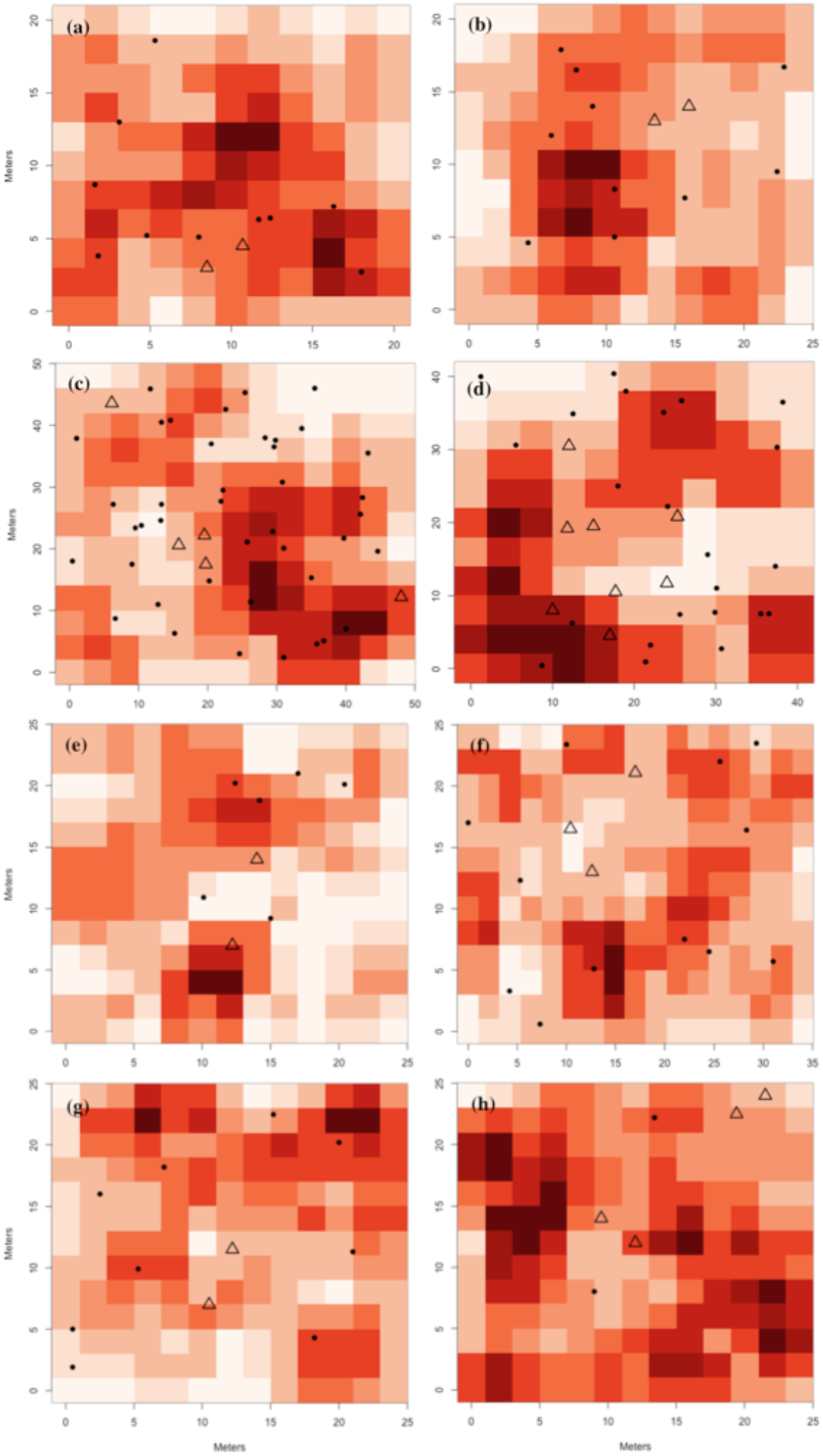
Graphical representations of ant species richness found at all plots. Points represent shade trees, triangles represent shade trees with a nest of *A. instabilis* and darker colour squares indicate more species. Graphs (a, b) represents high shade Plots H1 and H2; (c-f) represents moderate shade Plots M1-4; and (g, h) represent low shade Plots L1 and L2, respectively.

